# An IgG-Optimized Enzyme-Linked Lectin Assay (ELLA) for Quantitative Analysis of Immunoglobulin Glycosylation

**DOI:** 10.64898/2026.05.28.728458

**Authors:** Christine D. Wiggins, Douglas A. Lauffenbuger

## Abstract

Antibody Fc glycosylation is modulated in a variety of disease and immune response contexts, altering downstream functional responses including antibody-dependent cellular cytotoxicity through modified immune cell Fc receptor binding. Accessible, high-throughput glycosylation assays such as enzyme-linked lectin assays (ELLAs) are essential to advance understanding of glycosylation regulation and function. However, current ELLA protocols lack standardization and optimization, and results are reported out in arbitrary absorbance units, limiting reproducibility and cross-study comparability. We developed an optimized multi-lectin parallel ELLA with three specific improvements: systematic optimization of incubation times and reagent concentrations; incorporation of Protein A for IgG specificity; and use of commercially available bovine fetuin B as a quantitative surrogate standard for cross-study reproducibility. Our panel of 8 lectins, SNA, RCA, LCA, PHA-E, PHA-L, MAL-I, WGA, and DSL, cover the major IgG glycoforms. We demonstrate that our ELLA panel can reveal biologically relevant cytokine-induced plasticity of IgG glycosylation profiles in immortalized B cells.

## Introduction

Humoral immunity is an important arm of the immune response to infection or vaccination, in which antibodies specific to a particular antigen are produced at scale to enable the body to recognize that antigen and recruit immune cells to mount a response. Antibodies are comprised of a Fab portion, which binds to antigen, and an Fc region, which binds Fc receptors (FcRs) present on immune cells. The Fc region contains a conserved amino acid residue that allows for N-linked glycosylation, resulting in the addition of complex, branching glycan structures^1^.

Alterations in Fc glycosylation have been mechanistically linked to changes in immune cell Fc receptor (FcR) affinity and downstream effector functions, such as antibody-dependent cellular cytoxicity (ADCC) or induction of complement deposition^2,3^. These alterations have been observed across multiple inflammatory and autoimmune diseases^4–9^. Further, Fc glycosylation is dynamically modified during physiologic processes such as pregnancy and throughout the aging process^10–12^. Given the functional consequences of Fc N-glycosylation, reliable and scalable methods for quantitation are essential.

Two general strategies exist for measuring antibody glycosylation. In the first approach, glycans are enzymatically cleaved from the antibody and measured with analytical techniques such as liquid chromatography-mass spectrometry (LC-MS), matrix-assisted laser desorption/ionization time-of-flight (MALDI-TOF), or high-performance liquid chromatography (HPLC)^13–15^. These methods are high resolution, providing precise structural resolution and quantitation, but are expensive and require specialized equipment and expertise, reducing accessibility and scalability.

In the second approach, glycans are left intact on immunoglobulin and measured directly. This approach often leverages lectins, glycan-binding proteins with defined affinities for specific glycosylation patterns. Lectin-binding based approaches can be either chip-based or plate-based. In chip-based approaches, lectins are immobilized directly on microarrays, with the protein of interest then incubated across the array^16–18^. Quantification of binding of the protein of interest is then done via fluorescently labelled antibodies. In the plate-based approach, known as an enzyme-linked lectin assay (ELLA), the protein of interest is immobilized on a plate and probed with lectins conjugated to horseradish peroxidase, or another enzyme enabling colorimetric detection, and then quantified via colorimetric assay^19–22^. These lectin-based techniques are more accessible, require less specialized equipment and technical expertise, and enable higher-throughput analysis. Importantly, they measure glycans in their native protein context, capturing functional glycosylation motifs directly. Despite these advantages, lectin-based assays have not achieved standardized quantitative implementation.

Despite such advantages, lectin-based assays have not achieved standardized quantitative implementation. Published studies vary widely with respect to incubation times, reagent types and concentrations, coating strategies, and data processing methods. Additionally, there is a lack of reproducible calibration standards, which leads to reporting of results in arbitrary absorbance units. There is currently no reproducible calibration framework for ELLAs. This leads to limited quantitative comparability across experiments, between labs, or within the field at large. As a result, ELLA data are restricted to within-study relative comparisons.

We address these limitations through a two-pronged approach to ELLA improvement. First, we develop a harmonized, multi-lectin approach wherein we optimize key experimental parameters to allow for simple multi-lectin measurements, quantify technical variability, and establish a suggested workflow. We additionally develop and characterize a standardized calibration approach utilizing bovine fetuin B, a well-characterized, commercially available glycoprotein containing multiple diverse glycan structures that demonstrate high binding across multiple lectins, allowing for enhanced reproducibility and quantitative measurement. The incorporation of a fetuin-based standard curve can be replicated in ELLAs optimized for other glycoproteins, enabling this approach to be extended beyond immunoglobulin glycosylation. Finally, we utilize the improved ELLA workflow to investigate cytokine-driven glycosylation plasticity in an immortalized B cell culture model. Collectively, this work establishes a standardized and quantitatively calibrated framework for ELLA assays, enabling reproducible cross-study comparison of immunoglobulin glycosylation.

## Results

### Method overview

We established a set of reproducible ELLA assays optimized for detection of multiple glycoforms on immunoglobulin G, thereby enabling quantitative comparison of glycosylation across assays and studies. To this end, we selected a panel of eight lectins to capture the major forms of glycosylation on immunoglobulin produced by human cell lines as well as standard bioprocessing cell lines such as Chinese Hamster Ovary (CHO) cells (Table 1). These lectins have previously been shown to capture functional glycosylation motifs on immunoglobulin, and to bind to bovine fetuin B^22,23^. To enable quantitative comparison, we characterized a bovine fetuin B standard curve across all lectins. This standard runs alongside immunoglobulin samples, enabling colorimetric ELLA readouts to be translated into reproducible fetuin-equivalent units that can then be utilized in downstream analyses (Figure. 1A).

**Figure 1.**
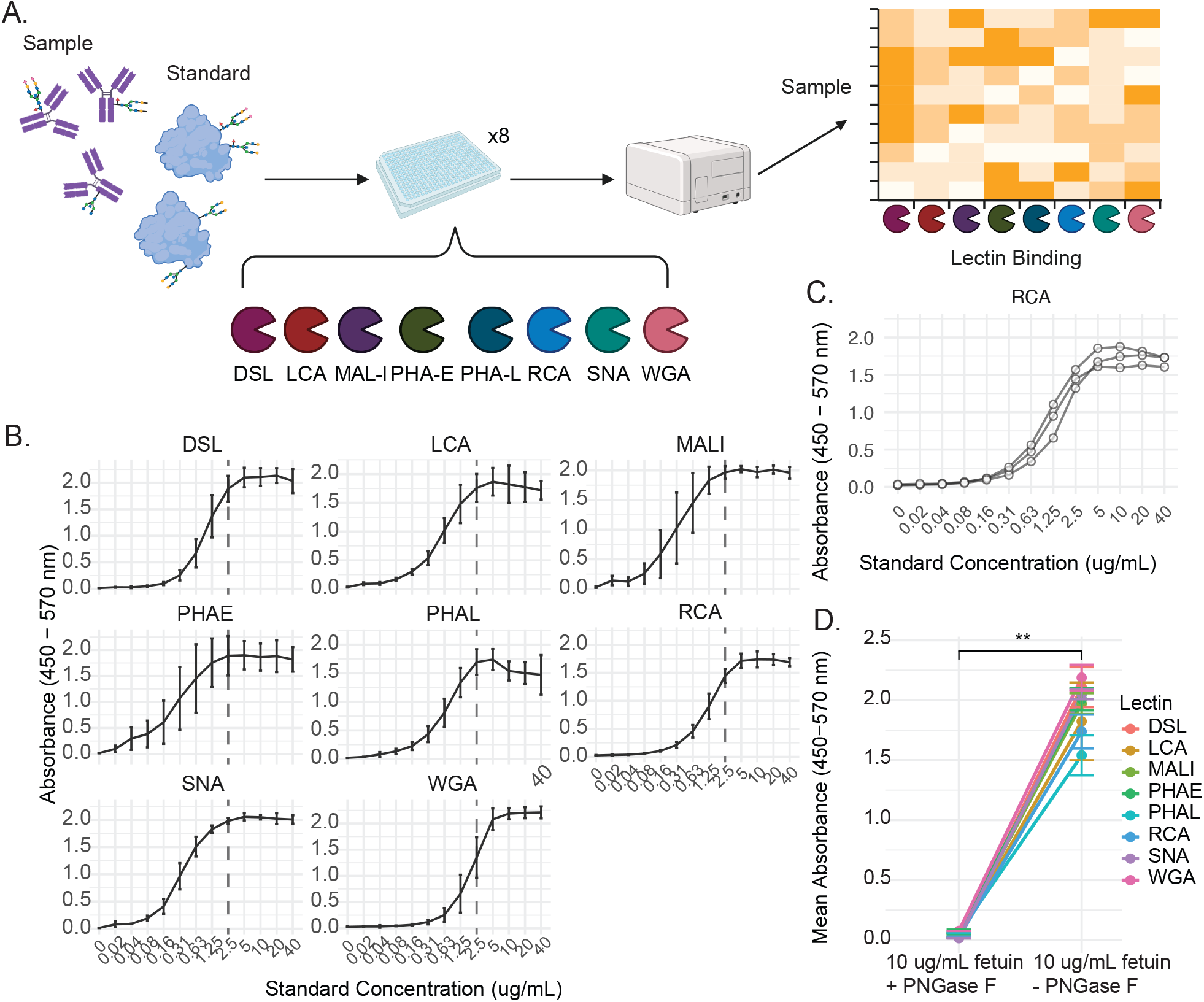
ELLA assay expanded to incorporate a bovine fetuin B standard curve. A) Schematic of workflow for ELLA process. B) Standard curves for serial dilutions of bovine fetuin B from 40 ug/mL to 0 ug/mL, for all 8 lectins. Each dot represents the mean of 3 experimental replicates, with error bars denoting one standard deviation from the mean. Each experimental replicate sample was measured in triplicate. C) Standard curves for three separate experiments measuring RCA binding to serial dilutions of bovine fetuin B from 40 ug/mL to 0 ug/mL. Each dot represents one experimental replicate, each of which was measured in triplicate. D) Mean absorbance for lectin binding to 10 ug/mL bovine fetuin B incubated with or without PNGase F to selectively cleave N-linked glycans. Each dot represents the mean of 3 experimental replicates, each measured in triplicate, for a given lectin. Error bars denote one standard deviation from the mean. Statistical significance is calculated with a paired t-test. Each lectin’s signal is reduced to background levels after deglycosylation with PNGase F, confirming N-glycan specific binding.

### Development of a Non-Proprietary Fetuin-Based Surrogate Standard Curve

To enable quantitative measurement of IgG glycosylation in cell culture supernatant via ELLA, a well-characterized standard is required. Standards should generally closely resemble the biomaterial tested, to ensure similar kinetics of binding and quantitative relevance. Ideally, a standard for this type of assay would be a highly glycosylated commercially available antibody. However, commercially available therapeutic antibodies such as rituximab do not exhibit all the forms of glycosylation necessary to make a good standard, as each antibody exhibits a distinct glycosylation profile. For example, rituximab lacks the sialic acid structures that bind SNA ^16,24^. Different therapeutic antibodies would thus be necessary to quantify each type of lectin-bound glycosylation. Meanwhile, bovine fetuin B is a well-characterized glycoprotein, with broad glycan diversity, commercially available, and non-proprietary, making it an ideal candidate for development of a repeatable ELLA standard. It has been previously utilized as a model glycoprotein in the development of many glycosylation measurement protocols. Further, the use of a surrogate standard, where a protein other than the one being assayed is used to generate a standard curve, has successfully been demonstrated in contexts such as ELISA measurement of toxins ^25^.

To define a dynamic range that worked across all 8 assays for ease of operation, we tested serial dilutions of bovine fetuin B in PBS from 40 ug/mL to 20 ng/mL for each lectin, across 3 separate assays (Figure 1B). As the upper limit of detection for most lectins began to plateau around 2.5 ug/mL as seen by the absorbance signal reaching maximum or near-maximum intensity, this was selected as the upper limit to the standard curves. Inter-assay variability was observed at all standard concentrations, consistent with known variability in colorimetric development due to variation in reagent incubation time, reagent temperature, mixing, and user variability (Figure 1C, Supplemental Figure 1), with a mean inter-assay coefficient of variance of 8.06%. To validate that signal observed from each lectin was coming from specific binding to fetuin N-glycosylation, we incubated bovine fetuin B with PNGase F, an enzyme that cleaves N-linked glycans. Across 3 separate experiments, cleavage of N-linked glycans abrogated lectin binding for all 8 lectins, confirming lectin signal dependence on the presence of N-linked glycosylation (Figure 1D).

### Systematic Optimization of Assay Parameters to Minimize Technical Variability

To minimize inter-assay variability and maximize dynamic range, we systematically optimized assay parameters such as lectin concentration and incubation. Previous work utilizing ELLAs typically employs a single concentration across multiple lectins, without optimization on a per-lectin basis. Here, we investigated three concentrations per lectin across the range of bovine fetuin B tested (Figure 2A). Lectins exhibit optimal results at different lectin concentrations. For most lectins, 1 ug/mL maximized dynamic range while maintaining low inter-assay variability (Table 2). SNA required a concentration of 0.1 ug/mL to prevent signal saturation, whereas LCA and MAL-I assays performed best at higher concentration of 10 ug/mL to achieve sufficient dynamic range. The effects of lectin concentration can be seen more finely by observing traces for each individual experiment, with lectins such as RCA exhibiting experimental consistency across all lectin concentrations, while experimental traces for SNA clearly reveal the enhanced experimental consistency at 0.1 ug/mL concentration (Supplemental Figure 1).

**Figure 2.**
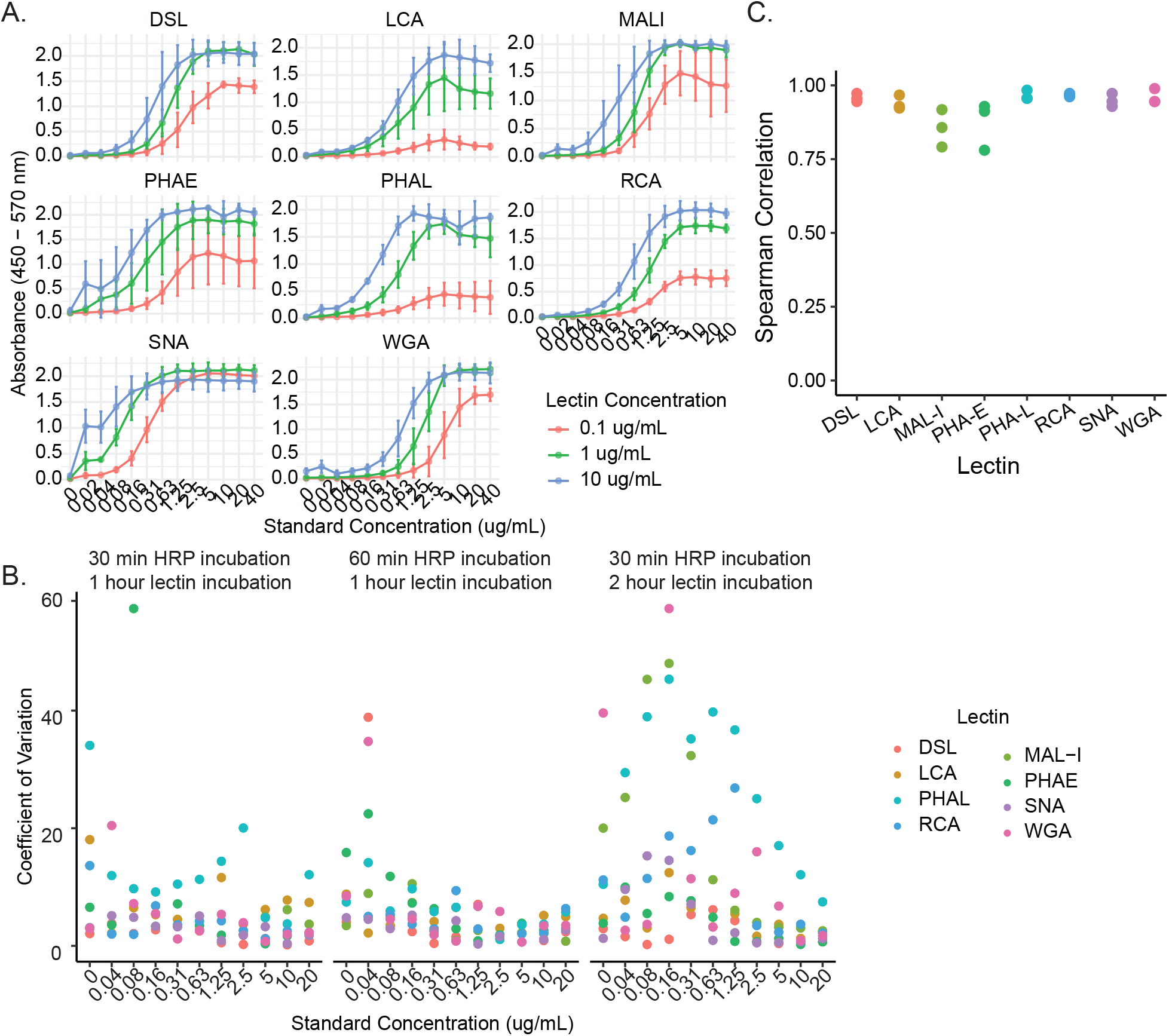
Optimizing lectin concentrations and incubation times improves ELLA performance. A) Standard curves for serial dilutions of bovine fetuin B from 40 ug/mL to 0 ug/mL, for all 8 lectins across 3 lectin concentrations. Each dot represents the mean of 3 experimental replicates, with error bars denoting one standard deviation from the mean. Each experimental replicate sample was measured in triplicate. B) Spearman correlations for each of the 3 pairwise comparisons of 3 experimental replicates per standard curve per lectin. C) Calculated coefficient of variance (CV) between three triplicate measurements per sample per lectin for each bovine fetuin B concentration in the standard curve. CVs are shown for (left to right) 30 minute HRP incubation with 1 hour lectin incubation, 60 minute HRP incubation with 1 hour lectin incubation, and 30 minute HRP incubation with 2 hour lectin incubation.

In addition to lectin concentration, lectin incubation time and horseradish peroxidase (HRP) incubation time also play a role in measurement consistency. To investigate this, we ran a truncated fetuin standard curve of 20 ug/mL to 40 ng/mL with absorbance measurements taken in triplicate. We characterized the coefficient of variance (CV) within these triplicate measurements for lectin incubation times, and then HRP incubation times (Figure 2B). The best results were observed with a 1 hour lectin incubation, with all CVs under 15% (1 hour: 5.8 +/-3.5%, 2 hour: 12 +/-10%) across all lectins. A 30-or 60-minute HRP incubation step was also tested, with comparable results (30 minute: 5.8 +/-3.5%, 60 minute: 5.2 +/-1.6%) and good average curves for both incubation times (Supplemental Figure 2A-C). Thus, we proceeded with a 30-minute HRP incubation period for experimental simplicity. To confirm experimental replicability, we computed Spearman correlations by lectin across 3 experimental replicates at the selected lectin concentrations and incubation times. We observed a high (>0.75 Spearman R) degree of reproducibility across the curve for each lectin, confirming the robust nature of the fetuin standard curve (Figure 2C).

### Integration of IgG Capture into a Quantitative Standardized Workflow

As common antibody-containing biofluids such as plasma or cell culture supernatants contain a variety of glycoproteins such as secreted cytokines in addition to the IgG we aim to focus on, we sought to incorporate and characterize a protein A (ProA) coating step within the assay to selectively capture IgG. Although ProA coating has previously been used in single-lectin ELLAs, comprehensive optimization across coating concentrations or utility in multiple lectin contexts has not been systematically evaluated^26^. We tested 3 concentrations of ProA loaded into each well, 2, 4, and 8 ug/mL. We then loaded a cell culture supernatant with a defined amount of human IgG at 9 serial dilutions, and measured galactosylation, initially just via RCA lectin binding. All ProA concentrations measured had comparable IgG capture as measured by RCA signal, for all sample concentrations measured (Figure 3A). Moving forward, a ProA concentration of 4 ug/mL was utilized. We next tested the efficacy of ProA in removing nonspecific signal from the supernatant. We utilized the same cell culture supernatant diluted to an IgG concentration of 6.5 ng/mL, the midpoint of our serial dilution curve, and either incubated the sample on untreated wells or on wells precoated with ProA. With direct binding to untreated wells, substantial signal was observed across all lectins measured, suggesting that the assay was detecting lectin binding to a variety of glycosylated supernatant proteins (Figure 3B). However, with an initial ProA capture step, signal was significantly reduced as compared to untreated wells (Wilcoxon signed-rank p = 0.0078) to close to what was observed in prior phosphate buffered saline (PBS) only samples. ProA coating is performed only for wells in which sample supernatants are loaded; fetuin standard curve wells remain uncoated to allow direct adsorption to the plate surface. We next examined the linearity of IgG dilution using two-fold dilutions of supernatant containing human IgG, incubated on ProA-coated wells alongside standard curves of bovine fetuin B (Figure 3C). Samples were measured in triplicate for all lectins. Across 7 of the 8 lectins, binding is detected at levels proportionate with sample concentration across all samples. For the 8^th^ lectin, PHA-L, which binds tri-antennary N-glycans, no detectable binding was observed across all supernatant concentrations. Many antibodies do not exhibit tri-antennary glycosylation, and the lack of binding of this lectin to our sample serves further evidence of lectin binding specificity^17^.

**Figure 3.**
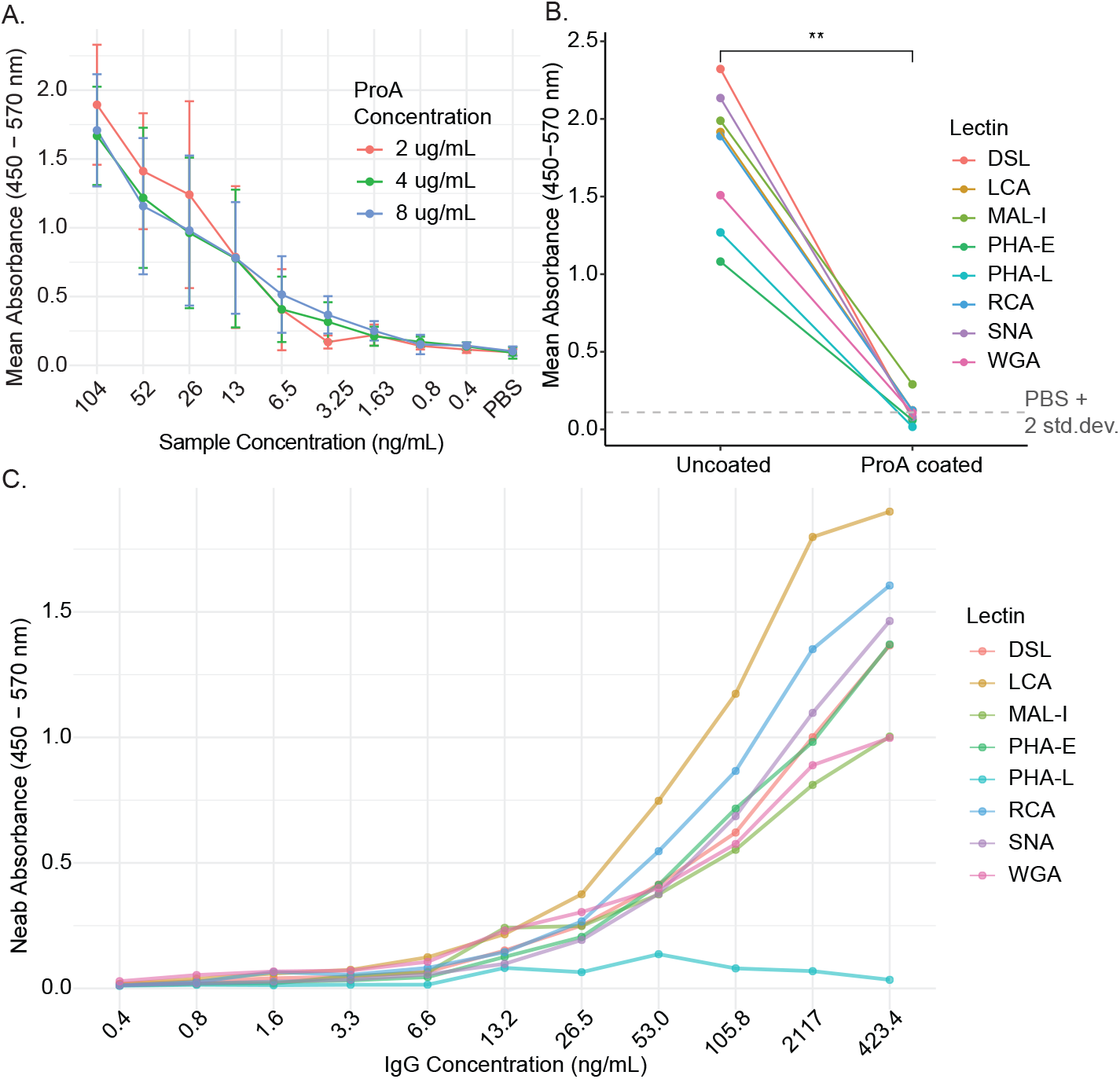
Incorporating Protein A capture step makes assay IgG-specific. A) Absorbance for serial dilution of a Bimm culture IgG-containing supernatant from 25x to 6400x, across 3 different ProA concentrations for plate coating. Dots represent the mean of 3 experimental replicates, each measured in triplicate. Error bars denote one standard deviation from the mean. All measurements performed with RCA lectin. B) Absorbance for 200x dilution of an IgG supernatant with either direct adsorption to the ELISA plate or incubation on a plate coated with 4 ug/mL ProA. Dots represent one experimental replicate, measured in triplicate, colored by lectin. Significance was calculated via paired t-test. C) Absorbance for serial dilution of an IgG-containing supernatant from 3x to 6400x, for all 8 lectins. Supernatant was incubated in plates coating with 4 ug/mL ProA. Dots represent one experimental replicate, measured in triplicate.

### Correlation between bovine fetuin B standard curve and IgG serial dilutions

To confirm the appropriateness of the surrogate standard, the serial dilution curves for both bovine fetuin B and supernatant IgG were compared for each lectin, plotting percentage of maximum observed absorbance as a function of protein concentration. The bovine fetuin B surrogate standard curve was similar to that using supernatant IgG, both of which were S-shaped curves well-fit using five-parameter logistic models (Figure 4A, Figure 4B). From these fits the protein concentrations of bovine fetuin B and supernatant IgG respectively at which the same percentage of maximum absorbance (20%-80%, at 10% intervals) was observed were calculated. For each of the 7 lectins with detectable binding to supernatant IgG, the relationship between bovine fetuin B concentration and supernatant IgG concentration at each absorbance percentage was linear, with R^2^ values of a linear regression ranging from 0.973 to 0.999 (Figure 4C). This excellent linearity demonstrates that bovine fetuin B is a suitable surrogate standard for a supernatant IgG ELLA, as both fetuin and IgG exhibit proportional lectin-binding kinetics across a relevant concentration range. All sample measurements on a given plate can thus be converted into fetuin equivalent values by way of the plate’s standard curve.

**Figure 4.**
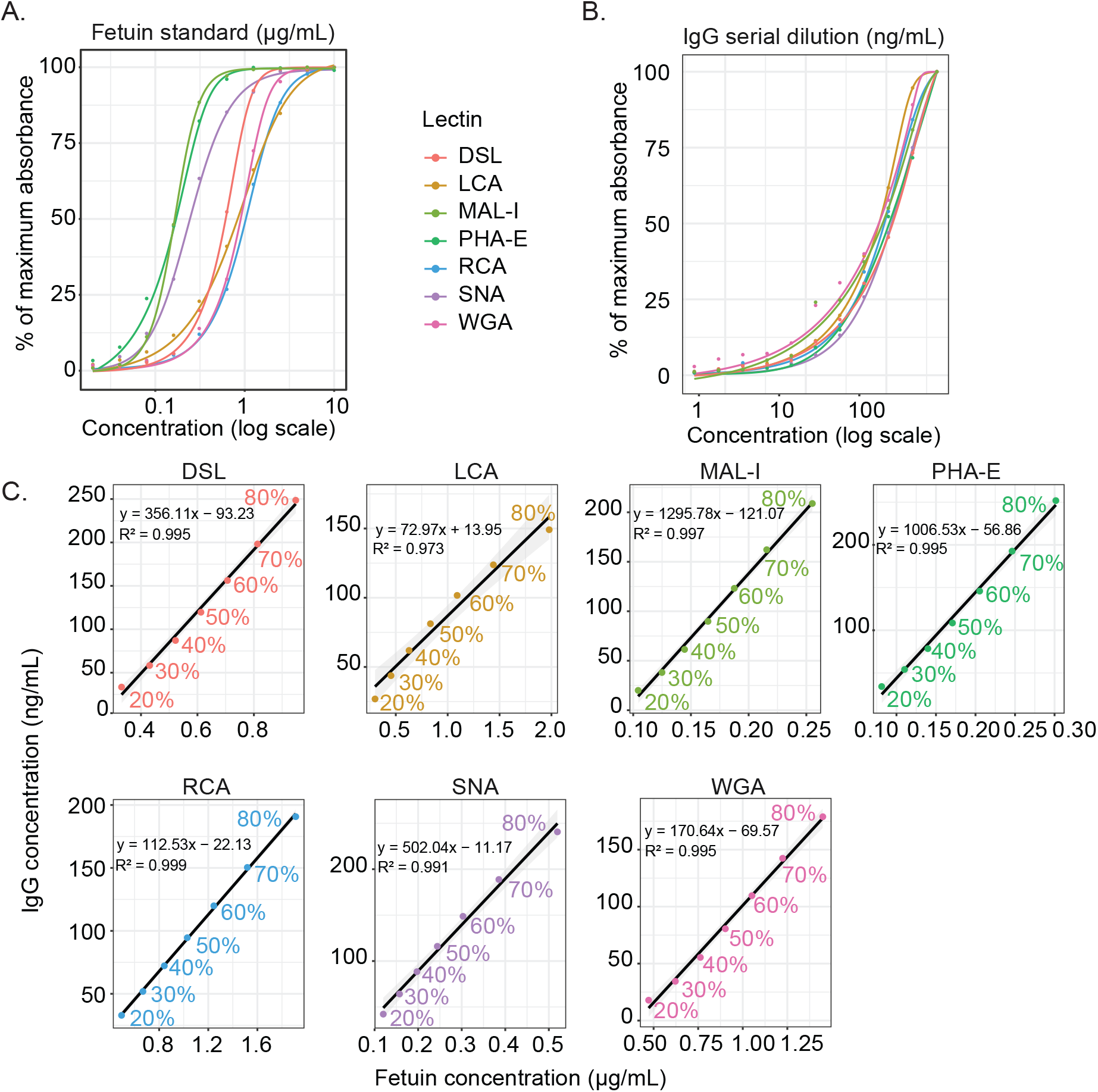
Bovine Fetuin B is an appropriate surrogate standard for IgG. A) Fetuin standard curves for all 7 IgG-binding lectins, with signal transformed to percentage of maximum background-corrected absorbance detected. Dots represent one experimental replicate, measured in triplicate. Curves were generated through 5 parameter logistic (5PL) fitted models. Run on the same plates simultaneously was B) IgG serial dilution curves for all 7 IgG-binding lectins, with signal transformed to percentage of maximum background-corrected absorbance detected. Dots represent one experimental replicate, measured in triplicate. Curves were generated through 5PL fitted models. C) 5PL-calculated IgG and bovine fetuin B concentrations at which a given percentage of maximum respective absorbance is estimated, per lectin. Linear regressions for each lectin individually are in black, with standard error in gray. Equations describe the linear regression fit for each lectin.

### Sensitivity of the Standardized ELLA to Detect Glycosylation Plasticity

The value of this assay is its ability to quickly quantify differences in glycosylation profiles between many samples. To illustrate this, we turned to immortalized B cells (“Bimms”) previously observed to exhibit cytokine stimuli-driven changes in secreted antibody glycosylation to test whether we could detect evidence of glycosylation plasticity in response to stimuli^27,28^. We utilized IFNg and IL-4 cytokine stimulation, either alone or pairwise, to determine whether antibody glycosylation produced under IFNg stimulation followed by IL-4 stimulation was different from antibody glycosylation produced under IL-4 stimulation alone, and similar. We chose immortalized B cells from two separate healthy donors, 4 unique clonal lines per donor to capture some of the diversity in clonal repertoire in humoral profiles. Clones were pooled per donor at equivalent cell numbers, 100,000 viable total cells per well in 12 well plates, to create Bimm pool 1 and Bimm pool 2. Cultures were grown on wells pre-coated with 200,000 irradiated IL-21-secreting YK6 cells, which express CD40L, as “feeder cells” to support Bimm growth^29^. Cultures were then stimulated with either 20 ng/mL IFNg or 50 ng/mL IL-4, with media replaced every 3 days. Conditions included stimulation with IL4 or IFNg for 6 days, IL-4 for 3 days followed by IFNg for 3 days (“IL-4_IFNg”), IFNg for 3 days followed by IL-4 for 3 days (“IFNg_IL-4”), or basal unstimulated media (Figure 5A). 4 biological replicates were cultured for each condition per Bimm pool, for a total of 40 samples.

**Figure 5.**
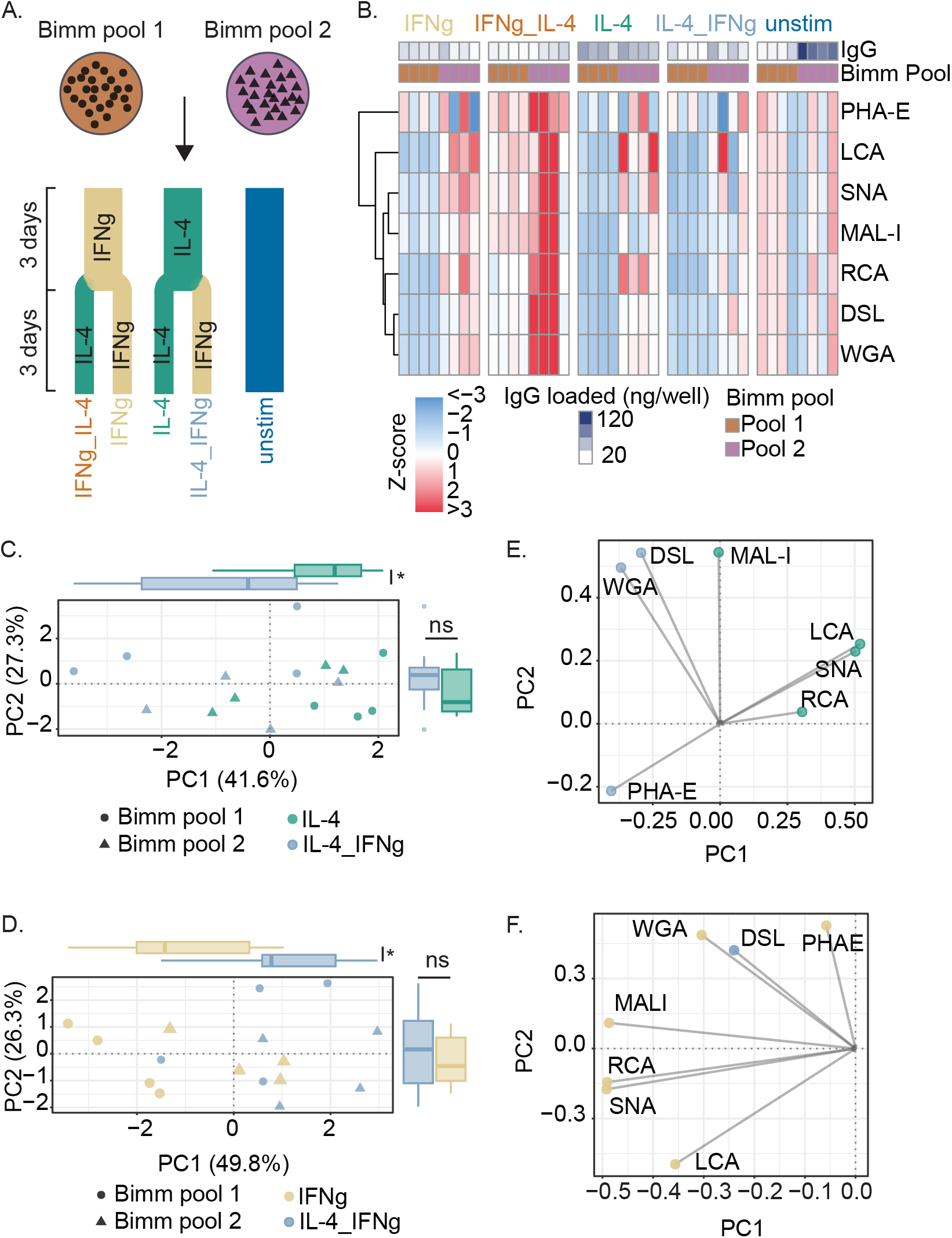
ELLAs can detect IgG glycosylation plasticity in an immortalized B cell system. A) Schematic of the experimental design for the glycosylation plasticity experiment. B) Heatmap of lectin-binding data per sample, z-scored via centering by the mean of a given Bimm pool and scaling by the standard deviation of that Bimm pool. Rows are clustered by Euclidean distance. Principal components analysis (PCA) scores plot of C) IL-4 and IL-4_IFNg stimulated samples, and D) IFNg and IL-4_IFNg stimulated samples. Boxplots on each axis represent the distribution of principal component (PC) 1 and PC2 scores per group, with statistical significance assessed via Mann-Whitney nonparametric test (* p < 0.05). PC labels represent the percentage of total lectin binding variance explained by each component. PCA loadings plots were generated for an E) IL-4 and IL-4_IFNg comparison, and an F) IFNg and IL-4_IFNg comparison. Loadings are in two dimensions, representing feature contribution to both PC1 and PC2. Lectin nodes are colored by which stimulation condition that lectin’s binding was univariately higher in.

Supernatant from each replicate was processed after 6 days of stimulation, with total IgG quantified via ELISA and glycosylation measured via 7 parallel ELLAs, excluding PHA-L. Total IgG produced differed across culture conditions, with IFNg_IL-4 exhibiting the lowest overall production (Supplemental Figure 3A). To normalize by Bimm pool, each lectin measurement was z-scored using the mean and standard deviation of the unstimulated samples from that Bimm pool. While differences in lectin binding between Bimm pools for a given stimulation condition were observed, overall IFNg_IL-4 samples showed higher binding across multiple lectins for both pools (Figure 5B). When compared to unstimulated samples, one of these lectins, MAL-I, binds at a univariately significantly higher level in IL-4 stimulated samples, while another, PHA-E, is significantly higher binding in IFNg_IL-4 samples (Supp Fig 3B), indicating stimulation conditions are inducing differential glycosylation. Though IFNg_IL-4 samples had the lowest overall supernatant IgG, this difference is not fully caused by remaining artifacts of differences in IgG loaded despite normalization, as Pearson correlations reveal a non-zero but weak negative correlation between lectin binding and titer across 6 of the 7 lectins (Supp Fig 3C).

Although most lectin measurements revealed subtle, non-significant differences between conditions, the knowledge that complex glycan structures are comprised of multiple glycan components led us to employ multivariate models to compare overall glycosylation profiles. Principal components analysis (PCA) was performed on lectin binding profiles for comparisons of interest, allowing for the reduction of 7-dimensional binding profiles to a smaller number of interpretable dimensions. When all stimulation conditions were analyzed together, IFNg_IL-4 separated from other groups (Supplemental Figure 3D), driven by elevated lectin binding across the entire panel (Supplemental Figure 3E). This unique profile indicates a highly glycosylated state that differs from samples exposed to IL-4 or IFNg alone, or both in the reverse sequential combination, suggesting that the order of cytokine-driven cellular signals fundamentally shapes the resultant glycosylation profile.

Pairwise PCAs revealed that neither the initial nor final stimulus alone set the glycosylation outcome. Comparing IL-4 to IL-4_IFNg, PC1 (capturing 41% of the variance in the relevant lectin binding data) differed significantly between the two conditions (Figure 5C). A loadings plot of lectin measurement contributions to PCA scores reveals that higher binding of LCA, SNA, and RCA, indicating higher levels of core fucose, terminal sialic acid, and galactose, are present in IL-4 treated samples. The stimulation with IFNg for the final 3 days appears to shift the profile away from this terminally decorated mature glycosylation profile. In a comparison to IFNg-only stimulated samples, IL-4_IFNg samples again exhibit a unique profile (Figure 5D). Differences in PC1 scores now appear to be driven by higher binding of most lectins, including LCA, SNA, RCA, DSL, WGA, and MAL-I to IFNg-only samples, indicating a highly glycosylated phenotype in IFNg-stimulated cells (Figure 5E). By contrast, cells pre-primed with IL-4 demonstrate a relatively more restrained IgG glycosylation profile.

## Discussion

IgG N-glycosylation modulates antibody Fc receptor binding and subsequent induction of downstream functional responses such as ADCC and complement activation^2^. Disease-specific dysregulation of glycosylation patterns has been characterized in autoimmune diseases as well as in immunomodulatory states such as pregnancy and aging, underscoring the clinical relevance of high-throughput, reliable characterization methods for immunoglobulin glycosylation.^4,5,10,12^.

These high-throughput assays are essential to enable large-scale studies leveraging multi-omics integration to identify glycosylation-driven immune phenotypes as well as to identify complex cellular and systems-level drivers of antibody glycosylation. Current glycan measurement approaches such as LC-MS enable detailed quantification of glycan species, but are often expensive and time consuming, and require access to specialized equipment and expertise, making them less than ideal for large-scale or exploratory studies. Meanwhile, lectin-based approaches such as ELLAs enable lower-resolution but higher throughput characterization of functional glycosylation structures in their native conformations.

Yet ELLA approaches lack standardization, with substantial variability in incubation times, lectin concentrations, coating strategies, and buffer usage, limiting reproducibility. Our systematic optimization demonstrates that lectin-specific concentration tuning can increase the dynamic range of the assay by up to two-fold while balancing inter-assay variability, with an overall average coefficient of variance of 8% achieved. Further, the incorporation of a bovine fetuin B surrogate standard curve addresses a critical gap, where ELLAs have typically reported binding results in arbitrary absorbance units, limiting cross-study comparisons. Bovine fetuin B’s non-proprietary, commercially available and inexpensive nature makes it an ideal candidate for use in these assays. After optimizing the incorporation of bovine fetuin B, we demonstrate its utility as a quantitative surrogate standard. We observe R^2^ > 0.97 correlation to IgG serial dilutions at proportionately equivalent signal levels across 7 IgG-relevant lectins, demonstrating that bovine fetuin B lectin-binding kinetics are substantially similar to those of IgG across our defined working ranges. Through this process, we enable reporting of ELLA results as fetuin-equivalent units, facilitating reproducible assays and multi-study data integration.

However, the use of bovine fetuin B has a few inherent limitations. Fetuin B’s 3 N-linked glycosylation sites are nearly always completely glycosylated with complex glycan structures, and rarely with high mannose glycans^30^. While fetuin works as a surrogate standard in an ELLA measuring IgG glycosylation as we have confirmed the binding of all IgG-relevant lectins of interest, if this method were extended to measure additional glycoproteins, or IgG produced in alternate culture systems that may produce different glycoforms, fetuin B would need to be validated for those applications.

Additionally, this approach could be extended beyond IgG with the incorporation of other types of capture molecules. While this protocol employs protein A to capture most IgG subclasses, other capture molecules such as isotype-specific antibodies could be employed to investigate glycosylation of IgM or IgA. However, an advantage of proA is its production in bacterial cells and thus lack of native glycosylation. Isotype-specific antibodies, or antibodies to other proteins of interest, are frequently produced in mammalian cells and thus are themselves glycosylated and would require further optimization, such as deglycosylation prior to use. In its current form, however, this approach could be used to investigate glycosylation of plasma IgG as well as purified therapeutic proteins with an IgG-like Fc domain, such as Fc-fusion proteins.

This eight lectin panel was chosen to cover major glycosylation features of IgG produced by human B cells and common bioprocessing cells such as CHO cells. We are not the first to employ this specific panel, which has been shown to provide good coverage of major relevant glycoforms and even enable further computational inference of more specific and fine-tuned glycan structures^22,31^. Within the panel, SNA and MAL-I detect a-2,6 and a-2,3 sialic acids, terminal structures whose presence has been observed to give IgG a more anti-inflammatory profile, while RCA binds to terminal galactose^32^. Detection of changes in these structures has been leveraged in diagnosis of and therapeutic design for autoimmune conditions such as Rheumatoid Arthritis ^11,33,34^. LCA detects core fucose, which has been targeted for removal in glycoengineering strategies to enhance the antibody-dependent cellular cytotoxicity-driven therapeutic effects of several approved monoclonal antibodies ^35–37^. DSL, WGA, and PHA-E detect glycan branching and antenna complexity^38^. Finally, PHA-L recognizes complex tri-antennary epitopes^38^. As human IgG predominantly lacks tri-antennary glycosylation structures, lack of PHA-L binding to our IgG samples supports the specificity of these lectin profiles^39^.

While this lectin panel captures many of the primary motifs of human and CHO-produced IgG glycosylation, end users may need to expand the panel if investigating either IgG produced by alternate expression systems or glycoproteins expressing different branching patterns.

To validate that our optimized ELLA panel and process can detect biologically relevant IgG glycosylation changes, we applied it to investigate glycosylation plasticity in immortalized B cell cultures sequentially stimulated with cytokines. Prior work has demonstrated that B cells respond to environmental stimuli including cytokines by producing immunoglobulin with differing glycosylation^23,40^, but the ability of B cells to rapidly respond to changes in microenvironment to pivot glycosylation responses is uncharacterized. Principal components analysis revealed that sequential cytokine stimulation produced glycosylation profiles distinct from either initial or final cytokine stimuli. IL-4 followed by IFNg stimulation looked neither like IL-4 stimulation nor IFNg stimulation alone, nor did it resemble reverse-order stimulation samples. This result demonstrates that our optimized ELLA panel and approach is sensitive enough to detect context-dependent glycosylation. Interestingly, differences in binding of individual lectins were not sufficient to discriminate between stimulation conditions, underscoring the added value of multivariate analysis combined with multi-lectin measurements to resolve nuanced shifts in glycosylation. This finding demonstrates the utility of parallel, multi-lectin measurements for detecting complex glycosylation shifts in biologically-relevant contexts.

## Methods and Materials

### Enzyme-linked lectin assays (ELLA)

Though variations on the ELLA protocol are described in this manuscript to characterize the effects of alterations of incubation times, standard concentrations, and other parameters, the final standardized method is as follows. 384-well Nunc MaxiSorp microtiter plates (ThermoFisher) were coated with 23 uL per well of 4 ug/mL Protein A (ProA, ThermoFisher) and incubated for 4 hours shaking at room temperature, with the exception of wells designated as standard wells. After 3 hours of incubation, bovine fetuin B (Sigma Aldrich, reconstituted at 1 mg/mL and stored at −20C) at a concentration of 2.5 ug/mL in PBS was prepared and serially diluted 1:2 to generate a nine-point standard curve, with a tenth standard curve point of PBS. This standard curve was added to the remaining wells, and the plate was left to finish the final hour of incubation. The plate was washed three times with a wash buffer of PBS and 0.2% Tween20 (PBST). We observed a substantial reduction in binding efficiency and increased well-to-well variability when PBST washes occurred prior to incubation with ProA or bovine fetuin B (data not shown). Thus, washes should occur only after all initial coating is complete. The plate was blocked with SynBlock blocking buffer (BIO-RAD) for 1 hour shaking at room temperature before another three washes with PBST. For biological measurements, 23 uL of media supernatant diluted 1:5 and 1:30 in PBS was added to ProA wells in triplicate, and PBS was added to standard wells to prevent drying. The plate was then incubated at least 12 hours overnight shaking at 4C. After a further 3 PBST washes, a solution of lectin (Vector Labs) in binding buffer (DPBS supplemented with 10 mM HEPES) was added to each well and an additional 1 hour shaking incubation at room temperature was performed. Lectin concentrations, sources, and specificities are noted in Table 1. Streptavidin-HRP (Strep B, R&D Systems) was added to all wells after a further 3 PBST washes, and the plate was left for 30 minutes shaking at room temperature. After 3 more PBST washes, ELISA TMB substrate (R&D Systems) was added to the plate and allowed to incubate until appropriate signal arose in the standard curve, typically 5-20 minutes. 2N sulfuric acid (R&D Systems) was then added to stop the reaction, and the plate was read via plate reader at 450 nm and 570 nm.

### ELLA Data Analysis

All data were adjusted by subtracting absorbance at 570 nm from absorbance measured at 450 nm. Any well with a 570 nm measurement over 0.15 was later inferred to have the maximum possible signal, as this threshold empirically correlated with visible precipitate formation, indicating excessive TMB substrate accumulation and over-saturation of signal. 5 parameter logistic models were then fit via the ‘nplr’ R package to fetuin standard curves after averaging triplicate measurements. After averaging triplicate measurements per sample, standard curves per plate were used to convert sample measurements to fetuin equivalents. In IgG lectin measurements other than dilution curves, IgG concentration loaded per well had to be between 7 and 150 ng/mL, a range observed to maintain linear response in ELLA characterization.

Absorbance measured had to be under the fetuin standard curve plateau absorbance, calculated as the maximum standard curve absorbance measurement minus 10% of the observed absorbance range over the entire curve. Fetuin equivalent measurements were log transformed from a tail-heavy distribution. Measurements were corrected for differences in IgG loaded per well by division by the log of IgG loaded per well. This correction yields a glycosylation measurement proportional to glycan density per unit IgG.

### Deglycosylation of Bovine Fetuin B

To demonstrate the specificity of lectin binding to N-glycans, Peptide-N-Glycosidase F (PNGase F) was used to cleave N-glycans from bovine fetuin B. 10 ug (10 uL of 1 mg/mL stock) bovine fetuin B was denatured via incubation with PNGase F (New England Biolabs) according to manufacturer protocol. Heat denaturation did not improve bovine fetuin B N-glycan cleavage efficiency (data not shown), so all generation of deglycosylated bovine fetuin B utilized native, non-denatured protein for protocol simplicity.

### Cell Culture and Stimulation

The human cell line “YK6” (YK6-CD40Lg-IL21) expressing CD40L and secreting IL-21 (a kind gift from the Hodson lab, University of Cambridge, generated as described previously^29^ was cultured in “YK6 media”: Advanced Roswell Park Memorial Institute (Advanced RPMI 1640, ThermoFisher) medium containing 10% fetal bovine serum (FBS, ThermoFisher) and 1% penicillin/streptomycin (ThermoFisher). YK6 cells were irradiated with 30 Gy via Gammacell Irradiator prior to freezing at 4e6 viable cells/mL (vc/mL) in freezing media (45 mL FBS, 5 mL dimethyl sulfoxide, Sigma Aldrich).

Influenza H7 HA antigen-specific immortalized B cell (Bimm) lines F2-13, F2-27, F2-29, F2-30, F5-1, F5-16, F5-17, F5-24 (all kind gifts from Dr. Kristin Boswell, Koup research group, Vaccine Research Center, generated as described previously^28^) were cultured in “Bimm media”: Iscove’s Modified Dulbecco’s Medium (IMDM, ThermoFisher) with 10% Ultra-low IgG FBS (ThermoFisher) and 1% penicillin/streptomycin (ThermoFisher). Irradiated YK6 cells were plated in YK6 media at 200,000 cells/well in 12 well tissue culture treated plates and allowed to adhere for 24 hours prior to addition of Bimms, which are then seeded at an initial density of 100,000 cells/well, split equally between clonal lines per donor (F2 lines, F5 lines) to create two distinct Bimm pools. Upon addition of Bimms, media was changed to the Bimm media as described. Cell counting of Bimms for seeding was done via hemocytometer on an EVOS M5000 imaging system using the green fluorescent channel to count only Bimm cells, which express green fluorescent protein (GFP), and not YK6 cells, which do not. Viable cell counts were determined using trypan blue exclusion. Bimms were grown for 6 days, with media changed at day 3. On day 6, media was replaced with fresh media supplemented with either 20 ng/mL recombinant human IFNg, 50 ng/mL recombinant human IL-4, or no cytokine (PeproTech). Concurrently, 200,000 fresh irradiated YK6 cells were added per well. Bimms were then grown in their initial stimulation media for 3 days. At day 9, media was replaced with fresh media and stimulations were switched where applicable in sequential stimulation conditions or maintained constant for single-stimulation or control conditions. After 3 additional days (12 total culture days), supernatant was harvested and frozen at −80C for future ELISA and ELLA analysis.

All cell lines used in this study were maintained in a humidified incubator (5% CO2) at 37C. All cell lines used in this study were confirmed free of mycoplasma bacteria via MycoAlert mycoplasma detection kit (Lonza).

### Antibody quantification via ELISA

Human total IgG ELISA kits (Invitrogen/Thermo Fisher Scientific) were adapted for use in a 384 well plate format and used to quantify total IgG in the supernatant according to manufacturer instructions. Supernatants were diluted 1:5, 1:20, 1:40, and 1:100, to fall within a linear dilution range. Standard curves were fit using five-parameter logistic (5PL) models using the ‘nplr’ package in R, and samples were quantified from the standard curve, with each sample measured in triplicate.

### Parallel ELLAs for culture supernatants

Supernatants were diluted 1:5 and 1:30, chosen based on measured total IgG range, and 7 parallel ELLAs were performed, with samples measured in triplicate. After data processing as described in section 4.2, for each sample-lectin pair the dilution (5x or 30x) was chosen on the following basis. IgG concentration loaded per well had to be between 7 and 150 ng/mL, a range observed to be within the linear range in ELLA characterization. Absorbance measured had to be under the fetuin standard curve plateau absorbance, calculated as the maximum standard curve absorbance measurement minus 10% of the observed absorbance range over the entire curve. If both dilutions passed both filters, the 30x dilution was preferentially chosen to minimize non-specific binding and potential saturation.

### Statistical analysis

All plots were visualized and analyzed using R (version 4.3.1). Unless otherwise noted, univariate statistical comparisons between measurements were assessed using a Mann-Whitney nonparametric test, with Benjamini-Hochberg multiple hypothesis test correction. When more than two groups were compared, Kruskal-Wallis tests were first utilized. Significant comparisons (p < 0.05) were then examined for pairwise significance via Mann-Whitney nonparametric tests. For PCA, IgG-normalized fetuin-equivalents were z-scored within lectin measurements (mean-centered and scaled to unit variance) to enable comparison of glycosylation across conditions.

Figures were edited in Adobe Illustrator (v2021) only for page positioning, color, and size to enable clear visualization.

## Acknowledgements

The authors thank Julia Zhong, Dr. Diana Gong, Dr. Anisha Datta, Dr. Laura Bahlmann, and Dr. Brian Joughin for their technical expertise and constructive discussions. We gratefully acknowledge the gift of the feeder cell line YK6-CD40Lg-IL21 from Dr. Daniel Hodson and Miriam Di Re, Hodson Lab, Wellcome-MRC Cambridge Stem Cell Institute, University of Cambridge. We additionally acknowledge the gift of the immortalized B cell lines from Dr. Kristin Boswell and Dr. Timothy Watkins, Koup Lab, Vaccine Research Center, National Institutes of Health.

This project has been funded in in part with federal funds from the National Institute of Allergy and Infectious Diseases, the NIH, and the Department of Health and Human Services under contract No. 75N93019C00071, grant No. AI181898, and grant No. U19AI135995. We would also like to thank the Fairbairn Family Fund for supporting this work, as well as Army ICB UARC Contract W911NF-19-D-0001. This work was further supported by a graduate student fellowship from the MIT-Takeda Fellow Program, as well as a graduate student fellowship from the Siebel Scholars Foundation. Several experimental schematics were created with BioRender.

## References

1. Wang, T. T. & Ravetch, J. V. Functional diversification of IgGs through Fc glycosylation. J. Clin. Invest. 129, 3492–3498 (2019).

2. Dekkers, G. et al. Decoding the Human Immunoglobulin G-Glycan Repertoire Reveals a Spectrum of Fc-Receptor- and Complement-Mediated-Effector Activities. Front. Immunol. 8, (2017).

3. Lofano, G. et al. Antigen-specific antibody Fc glycosylation enhances humoral immunity via the recruitment of complement. Sci. Immunol. 3, eaat7796 (2018).

4. Deng, X. et al. Changes of serum IgG glycosylation patterns in rheumatoid arthritis. Clin. Proteomics 20, 7 (2023).

5. Azzam, T. et al. Asymmetrically glycosylated IgG1 antibodies are universal and drive human disease. Nat. Commun. 17, 383 (2025).

6. de Haan, N., Falck, D. & Wuhrer, M. Monitoring of immunoglobulin N- and O-glycosylation in health and disease. Glycobiology 30, 226–240 (2019).

7. Haslund-Gourley, B. S. et al. Host glycosylation of immunoglobulins impairs the immune response to acute Lyme disease. eBioMedicine 100, (2024).

8. Kennedy, P. G. et al. Aberrant Immunoglobulin G glycosylation in Multiple Sclerosis. J. Neuroimmune Pharmacol. Off. J. Soc. NeuroImmune Pharmacol. 17, 218–227 (2022).

9. Irvine, E. B. & Alter, G. Understanding the role of antibody glycosylation through the lens of severe viral and bacterial diseases. Glycobiology 30, 241–253 (2020).

10. Bondt, A. et al. Immunoglobulin G (IgG) Fab Glycosylation Analysis Using a New Mass Spectrometric High-throughput Profiling Method Reveals Pregnancy-associated Changes. Mol. Cell. Proteomics MCP 13, 3029–3039 (2014).

11. van de Geijn, F. E. et al. Immunoglobulin G galactosylation and sialylation are associated with pregnancy-induced improvement of rheumatoid arthritis and the postpartum flare: results from a large prospective cohort study. Arthritis Res. Ther. 11, R193 (2009).

12. Frkatović-Hodžić, A. et al. Mapping of the gene network that regulates glycan clock of ageing. Aging 15, 14509–14552 (2023).

13. Huffman, J. E. et al. Comparative Performance of Four Methods for High-throughput Glycosylation Analysis of Immunoglobulin G in Genetic and Epidemiological Research*. Mol. Cell. Proteomics 13, 1598–1610 (2014).

14. Kosuge, H. et al. Highly sensitive HPLC analysis and biophysical characterization of N-glycans of IgG-Fc domain in comparison between CHO and 293 cells using FcγRIIIa ligand. Biotechnol. Prog. 36, e3016 (2020).

15. Jansen, B. C. et al. MALDI-TOF-MS reveals differential N-linked plasma- and IgG-glycosylation profiles between mothers and their newborns. Sci. Rep. 6, 34001 (2016).

16. Zhang, L., Luo, S. & Zhang, B. The use of lectin microarray for assessing glycosylation of therapeutic proteins. mAbs 8, 524–535 (2016).

17. Luo, S. & Zhang, B. A tailored lectin microarray for rapid glycan profiling of therapeutic monoclonal antibodies. mAbs 16, 2304268 (2024).

18. Tsui, C. K. et al. CRISPR screens and lectin microarrays identify high mannose N-glycan regulators. Nat. Commun. 15, 9970 (2024).

19. Thompson, R., Creavin, A., O’Connell, M., O’Connor, B. & Clarke, P. Optimization of the enzyme-linked lectin assay for enhanced glycoprotein and glycoconjugate analysis. Anal. Biochem. 413, 114–122 (2011).

20. Klukova, L. et al. Glycoprofiling as a novel tool in serological assays of systemic sclerosis: A comparative study with three bioanalytical methods. Anal. Chim. Acta 853, 555–562 (2015).

21. Srinivasan, K. et al. A Quantitative Microtiter Assay for Sialylated Glycoform Analyses Using Lectin Complexes. J. Biomol. Screen. 20, 768–778 (2015).

22. Li, H. et al. LeGenD: determining N-glycoprofiles using an explainable AI-leveraged model with lectin profiling. BioRxiv Prepr. Serv. Biol. 2024.03.27.587044 (2024) doi:10.1101/2024.03.27.587044.

23. Cao, Y. et al. Cytokines in the Immune Microenvironment Change the Glycosylation of IgG by Regulating Intracellular Glycosyltransferases. Front. Immunol. 12, 724379 (2022).

24. Huang, W., Giddens, J., Fan, S.-Q., Toonstra, C. & Wang, L.-X. Chemoenzymatic Glycoengineering of Intact IgG Antibodies for Gain of Functions. J. Am. Chem. Soc. 134, 12308–12318 (2012).

25. Wang, Y., Li, P., Zhang, Q., Hu, X. & Zhang, W. A toxin-free enzyme-linked immunosorbent assay for the analysis of aflatoxins based on a VHH surrogate standard. Anal. Bioanal. Chem. 408, 6019–6026 (2016).

26. Klukova, L. et al. Glycoprofiling as a novel tool in serological assays of systemic sclerosis: A comparative study with three bioanalytical methods. Anal. Chim. Acta 853, 555–562 (2015).

27. Wiggins, C. D. et al. Multivariate analysis of glycogenes reveals coordinated regulation of immunoglobulin glycosylation in an immortalized human B cell system. 2025.09.30.679651 Preprint at 10.1101/2025.09.30.679651 (2025).

28. Boswell, K. L. et al. Application of B cell immortalization for the isolation of antibodies and B cell clones from vaccine and infection settings. Front. Immunol. 13, (2022).

29. Caeser, R., Gao, J., Di Re, M., Gong, C. & Hodson, D. J. Genetic manipulation and immortalized culture of ex vivo primary human germinal center B cells. Nat. Protoc. 16, 2499–2519 (2021).

30. Cao, L. et al. Global site-specific N-glycosylation analysis of HIV envelope glycoprotein. Nat. Commun. 8, 14954 (2017).

31. Yom, A. J., Chiang, A. W. T. & Lewis, N. E. A Boltzmann Model Predicts Glycan Structures from Lectin Binding. http://biorxiv.org/lookup/doi/10.1101/2023.06.03.543532 (2023) doi:10.1101/2023.06.03.543532.

32. Bojar, D. et al. A Useful Guide to Lectin Binding: Machine-Learning Directed Annotation of 57 Unique Lectin Specificities. ACS Chem. Biol. 17, 2993–3012 (2022).

33. Kaneko, Y., Nimmerjahn, F. & Ravetch, J. V. Anti-Inflammatory Activity of Immunoglobulin G Resulting from Fc Sialylation. Science 313, 670–673 (2006).

34. Magorivska, I. et al. Glycosylation of random IgG distinguishes seropositive and seronegative rheumatoid arthritis. Autoimmunity 51, 111–117 (2018).

35. Chiang, A. W. et al. Modulating carbohydrate–protein interactions through glycoengineering of monoclonal antibodies to impact cancer physiology. Curr. Opin. Struct. Biol. 40, 104–111 (2016).

36. Yamane-Ohnuki, N. et al. Establishment of FUT8 knockout Chinese hamster ovary cells: an ideal host cell line for producing completely defucosylated antibodies with enhanced antibody-dependent cellular cytotoxicity. Biotechnol. Bioeng. 87, 614–622 (2004).

37. Shields, R. L. et al. Lack of Fucose on Human IgG1 N-Linked Oligosaccharide Improves Binding to Human FcγRIII and Antibody-dependent Cellular Toxicity*. J. Biol. Chem. 277, 26733–26740 (2002).

38. Bojar, D. et al. A Useful Guide to Lectin Binding: Machine-Learning Directed Annotation of 57 Unique Lectin Specificities. ACS Chem. Biol. 17, 2993–3012 (2022).

39. Huhn, C., Selman, M. H. J., Ruhaak, L. R., Deelder, A. M. & Wuhrer, M. IgG glycosylation analysis. PROTEOMICS 9, 882–913 (2009).

40. Wang, J. et al. Fc-Glycosylation of IgG1 is Modulated by B-cell Stimuli. Mol. Cell. Proteomics MCP 10, M110.004655 (2011).

