## Supplemental Figures and Tables for "An IgG-Optimized Enzyme-Linked Lectin Assay (ELLA) for Quantitative Analysis of Immunoglobulin Glycosylation"

**Table 1.** Lectins, specificities, sources, and final concentrations for optimized ELLA assay.

| Lectin | Glycan target | Source, Catalog Number | Concentration for ELLA ($\mu$g/mL) |
|---|---|---|---|
| Sambucus Nigra Agglutinin (SNA) | $\alpha$2-6-sialylated LacNAc | Vector Laboratories, B-1305-2 | 0.1 |
| Ricinus Communis Agglutinin I (RCA) | Terminal Type 2 LacNAc | Vector Laboratories, B-1085-1 | 1 |
| Lens Culinaris Agglutinin (LCA) | Core fucose | Vector Laboratories | 10 |
| Maackia Amurensis-I (MAL-I) | $\alpha$2-3-sialylated LacNAc | Vector Laboratories, B-1315-2 | 10 |
| Phaseolus Vulgaris Erythroagglutinin (PHA-E) | Bisecting GlcNAc with Type 2 LacNAc | Vector Laboratories, B-1125-2 | 1 |
| Wheat Germ Agglutinin (WGA) | Terminal GlcNAc$\beta$<br>Terminal GlcNAc$\alpha$<br>Terminal NAc containing glycans | Vector Laboratories, B-1025-5 | 1 |
| Datura Stramonium Lectin (DSL) | Chitin<br>Type 2 poly LacNAc | Vector Laboratories, B-1185-2 | 1 |
| Phaseolus Vulgaris Leucoagglutinin (PHA-L) | $\beta$1-6 branched N-glycans | Vector Laboratories, B-1115-2 | 1 |

**Table 2.** Concentration-specific lectin performance metrics.

| Lectin | Lectin concentration (ug/mL) | Mean coefficient of variance (%) | Mean dynamic range (absorbance units) |
|---|---|---|---|
| DSL | 0.1 | 6.1 | 0.96 |
| DSL | 1.0 | 7.9 | 1.86 |
| DSL | 10.0 | 5.4 | 1.96 |
| LCA | 0.1 | 6.5 | 0.24 |
| LCA | 1.0 | 9.1 | 1.29 |
| LCA | 10.0 | 5.7 | 1.68 |
| MAL-I | 0.1 | 9.9 | 1.26 |
| MAL-I | 1.0 | 8.5 | 1.91 |
| MAL-I | 10.0 | 10.4 | 1.86 |
| PHA-E | 0.1 | 11.8 | 1.13 |
| PHA-E | 1.0 | 8.7 | 1.79 |
| PHA-E | 10.0 | 10.7 | 1.64 |
| PHA-L | 0.1 | 9.9 | 0.37 |
| PHA-L | 1.0 | 7.4 | 1.66 |
| PHA-L | 10.0 | 10.4 | 1.81 |
| RCA | 0.1 | 6.4 | 0.57 |
| RCA | 1.0 | 6.4 | 1.41 |
| RCA | 10.0 | 8.7 | 1.85 |
| SNA | 0.1 | 9.2 | 1.92 |
| SNA | 1.0 | 5.1 | 1.81 |
| SNA | 10.0 | 4.7 | 0.93 |
| WGA | 0.1 | 9.9 | 0.34 |
| WGA | 1.0 | 8.8 | 1.32 |
| WGA | 10.0 | 10.7 | 1.85 |

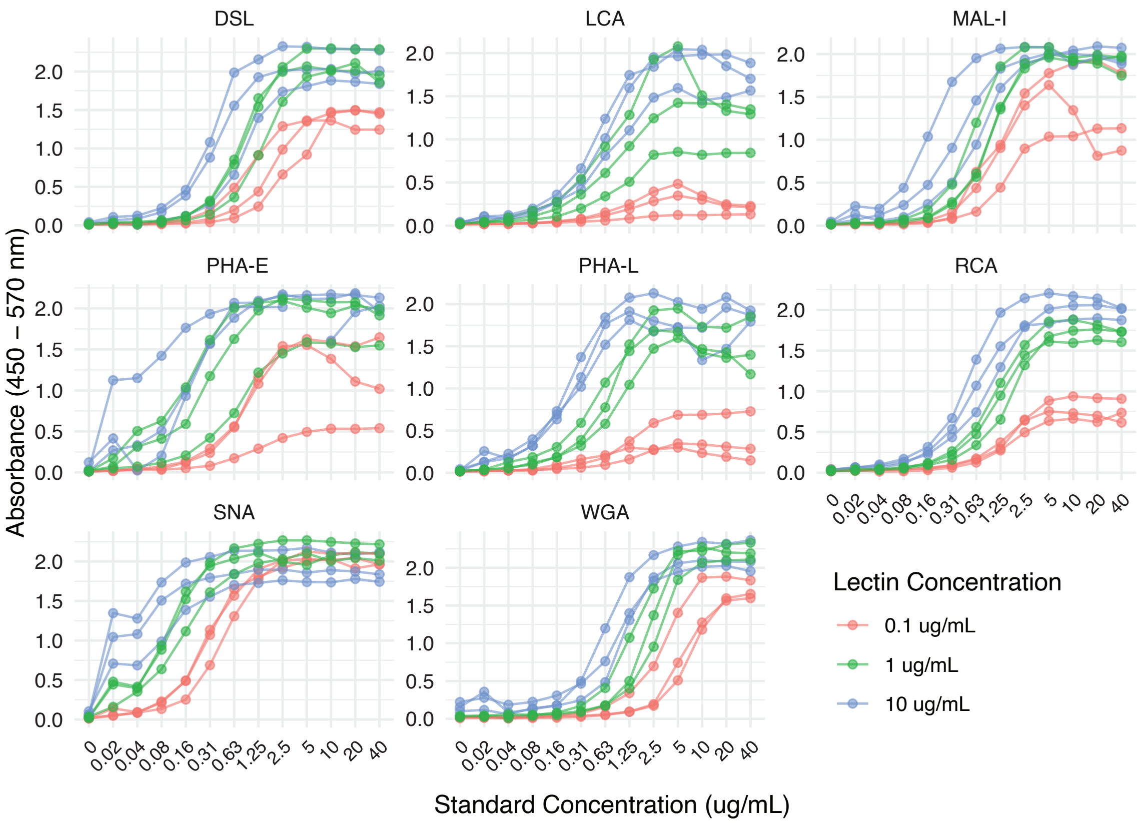


**Supplemental Figure 1. Selected lectin concentrations exhibit run-to-run consistency.** A) Standard curves for serial dilutions of bovine fetuin B from 40 ug/mL to 0 ug/mL for all 8 lectins at 3 different concentrations each. Each dot represents one experimental replicate, each of which was measured in triplicate.

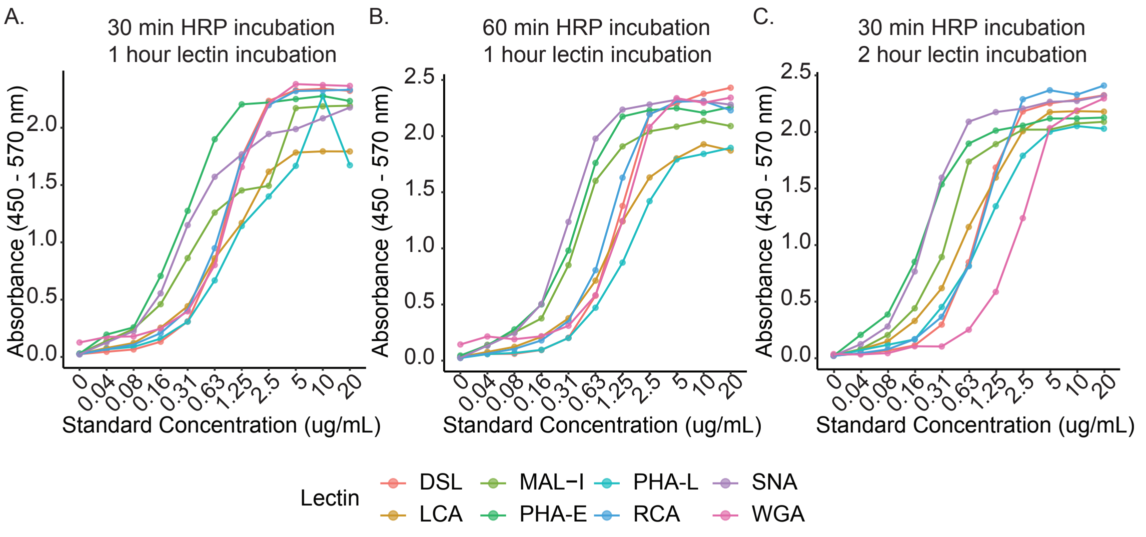


**Supplemental Figure 2. Lectin-specific standard curves vary with HRP and lectin incubation times.** Standard curves for serial dilutions of bovine fetuin B from 20 ug/mL to 0 ug/mL for all 8 lectins with A) a 30 minute HRP incubation and 1 hour lectin incubation, B) a 60 minute HRP incubation and 1 hour lectin incubation, and C) a 30 minute HRP incubation and 2 hour lectin incubation. Samples were measured in triplicate.

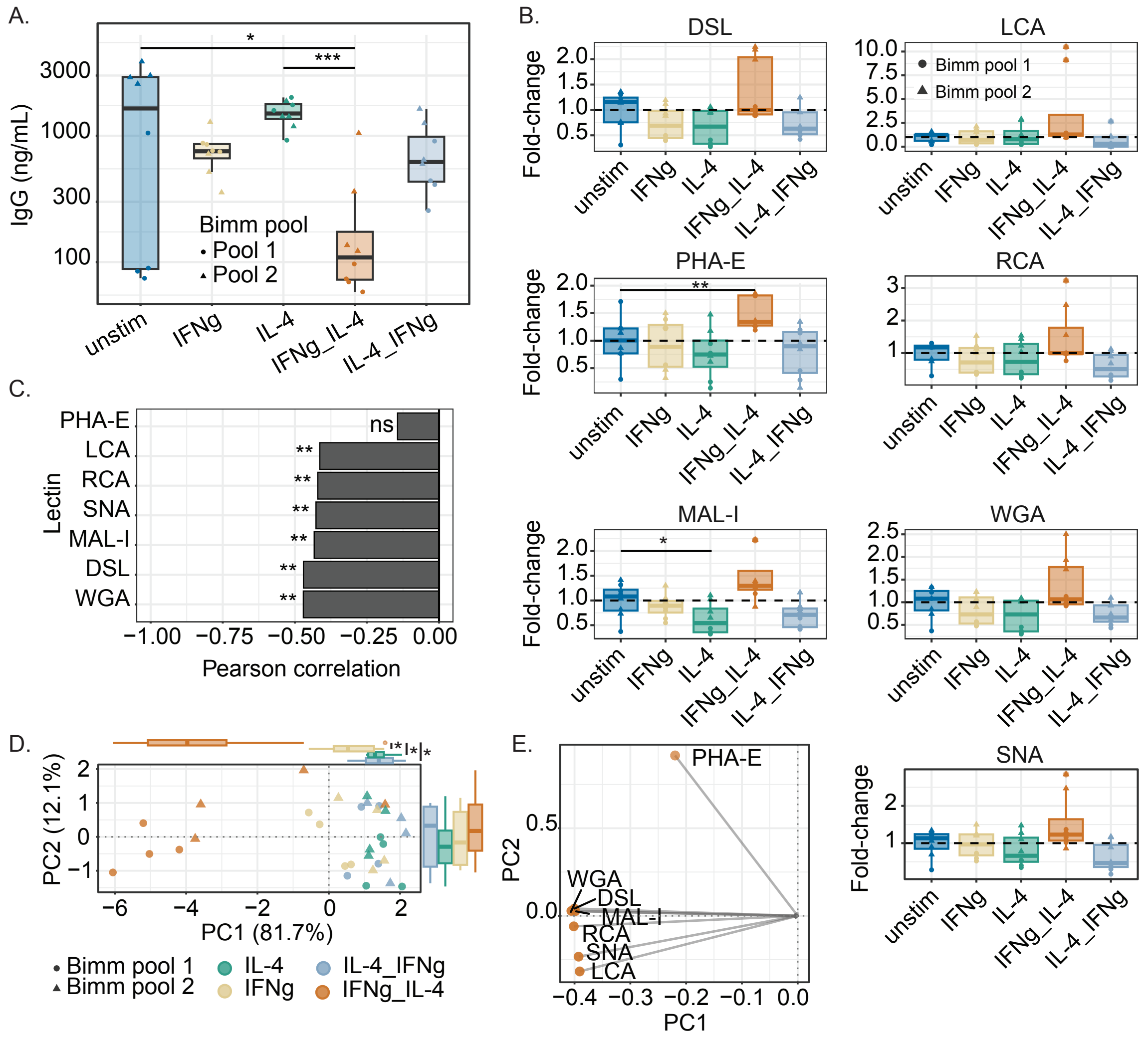


**Supplemental Figure 3. Lectin binding reveals glycosylation differences between cytokine stimulation conditions.** A) Total IgG in all supernatants was measured via ELISA, with differences between conditions tested with Kruskal-Wallis test followed by Dunn's post-hoc test, with Bonferroni correction. B) Fold-change of each measurement to pool-specific unstimulated mean value for each lectin. Dashed line indicates a fold-change of 1, where a sample would be equivalent to the mean value. Boxplots display the median value, with lower and upper hinges representing the 25th and 75th percentiles, and whiskers plotted from the hinges to the largest and smallest values at most 1.5x the interquartile range. Statistical significance between groups for each lectin was calculated with Kruskal-Wallis tests, with significant lectins then tested pairwise between unstimulated and stimulated conditions with Mann-Whitney nonparametric tests, multiple hypothesis corrected with Benjamini-Hochberg. C) Pearson correlation values between lectin binding values and IgG loaded per well for ELLA assays. Statistical significance indicates that a correlation is significantly different from zero, though all correlations fall below 0.5, indicating a weak relationship. D) Principal components analysis (PCA) scores plot of IL-4, IFNg, IFNg_IL-4, and IL-4_IFNg stimulated samples. Boxplots on each axis represent the distribution of principal component (PC) 1 and PC2 scores per group, with statistical significance assessed via Kruskal-Wallis, with significantly different PCs then tested pairwise with Mann-Whitney nonparametric tests (* $p < 0.05$). PC labels represent the percentage of total lectin binding variance explained by each component. E) PCA loadings plot for the IL-4, IFNg, IFNg_IL-4 and IL-4_IFNg comparison. Loadings are in two dimensions, representing feature contribution to both PC1 and PC2. Lectin nodes are colored by which stimulation condition that lectin's binding was univariately higher in.
